# Integrating conventional tagging and acoustic telemetry improves estimates of post-release survival in a highly targeted reef fish

**DOI:** 10.64898/2026.03.16.711647

**Authors:** A. Challen Hyman, Angela Collins, Chloe Ramsay, Sean Wilms, Micheal S. Allen, Luiz Barbieri, Thomas K. Frazer

## Abstract

Accurate estimation of post-release survival is fundamental to fisheries stock assessment and effective management. Conventional tag-return studies and acoustic telemetry are commonly used to estimate this probability, yet each approach has limitations when applied independently. Using gag (*Mycteroperca microlepis*) as a case study, we integrated data from a large-scale conventional tagging program and an acoustic telemetry experiment within a discrete-time statistical modeling framework that links relative recapture risk with telemetry-derived fate. This approach enabled estimation of post-release survival across a broad gradient of capture depths representative of recreational fishing conditions. Estimated survival was high in shallow waters (≈97%) but declined with increasing capture depth, consistent with depth-related barotrauma. Applying model predictions to depth distributions from the recreational fishery yielded annual and monthly post-release survival probabilities. Annual estimates were consistent with values assumed in recent stock assessments, while monthly values highlighted seasonal patterns potentially relevant for management. This integrated framework advances post-release survival estimation by combining the extensive sample sizes and environmental coverage characteristic of conventional tagging data with the direct fate observations provided by acoustic telemetry, and offers a transferable approach for other highly targeted fisheries.

## Introduction

Accurate estimation of recreational post-release survival is critical for understanding the population-scale consequences of fishing practices. Marine fisheries in industrialized nations are increasingly dominated by the recreational sector (Ihde et al. 2011; Hyder et al. 2018; Shertzer et al. 2019; Brownscombe et al. 2019; Bower et al. 2020), with recreational harvest in some regions routinely exceeding commercial catch (e.g., Shertzer et al. 2019). For many highly targeted and intensively managed species, regulatory discards may substantially outnumber retained harvest (Runde et al. 2019; Shertzer et al. 2019; Shertzer et al. 2024). Under these conditions, low post-release survival of discarded fish can contribute disproportionately to total fishing mortality, potentially undermining management objectives and conservation targets (Coggins et al. 2007; Tetzlaff et al. 2013; Shertzer et al. 2024).

Two common approaches for estimating post-release survival in recreational fisheries are conventional tagging and acoustic telemetry, both of which allow researchers to monitor the fate of released fish over time. In conventional tagging studies, fish are captured, fitted with external tags, and released back into the wild (e.g., Rudershausen et al. 2014; Sauls 2014; Runde et al. 2019; Stallings et al. 2023). Tags typically display a unique identification number, contact information, and a reward notice to encourage reporting by anglers. Upon recapture, tag returns provide information on the timing and location of recovery, enabling inference of survival over the intervening period. Data collected at the time of tagging – such as capture date, location, size, condition, and environmental variables – can be used to assess factors influencing recapture rates, which are interpreted as an index of relative post-release survival. Acoustic telemetry studies employ electronic tags that emit coded acoustic signals detected by stationary receiver arrays (e.g., Curtis et al. 2015; Bohaboy et al. 2020; Rudershausen et al. 2025; Zimmermann et al. 2025). This technology allows near-continuous monitoring of individual fish, providing high-resolution data on movement patterns and timing of mortality events.

While both approaches provide valuable insights, they differ in spatial extent, sample size, and the type of survival inference they support. Conventional tagging programs can cover large areas and release hundreds to thousands of fish over broad temporal and environmental gradients (e.g., Sauls 2014; Rudershausen et al. 2019; Rudershausen et al. 2023; Hyman et al. 2026b), but reporting rates are generally unknown. As a result, only *relative* survival can typically be estimated using recapture ratios (i.e., “relative risk”; Hueter et al. 2006; Rudershausen et al. 2014; Sauls 2014; Runde et al. 2019; Rudershausen et al. 2020), with absolute survival inferred only under specific conditions or from external data. In contrast, acoustic telemetry facilitates estimation of absolute post-release survival based on telemetry-derived fate assignment (e.g., Runde et al. 2021), yet these studies are generally constrained in spatial extent and sample size due to logistical and cost limitations (e.g., Alonso-Fernández et al. 2022; Bohaboy et al. 2020; Runde et al. 2021; Rudershausen et al. 2025; Zimmermann et al. 2025).

One approach to overcome these inherent limitations is to combine multiple methodologies. Electronic tagging data can provide estimates of absolute post-release survival for fish under specific release conditions, including size, handling stress, and environmental context (e.g., Rudershausen et al. 2019; Alonso-Fernández et al. 2022). These high-resolution data allow researchers to directly observe mortality events, providing a benchmark or “anchor” for survival estimates. When paired with large-scale conventional tagging data, relative survival among fish from different release environments can be scaled to absolute values. This integrative framework allows extrapolation of absolute post-release survival across broader environmental gradients and fish populations than either method alone. Despite the complementary nature of these approaches, few studies have formally integrated conventional tag-return and acoustic telemetry data within a unified statistical framework (Pollock et al. 2004; Bacheler et al. 2009; Benoît et al. 2020; Scheffel et al. 2020), and none – to the best of our knowledge – have applied such models to estimate absolute post-release survival from observed data.

In this study, we constructed and applied a discrete-time statistical model to conventionally and acoustically tagged gag (*Mycteroperca microlepis*) captured and released offshore of Tampa Bay, USA. This approach facilitated extrapolation of absolute post-release survival across broader environmental gradients, producing more robust, management-relevant estimates than either method alone. Our study demonstrates how integrating multiple datasets can improve understanding of post-release survival patterns, inform stock assessments, and guide evidence-based fisheries management.

## Methods

We analyzed tag-return records from 3,363 gag collected between 2009 and 2012 as part of the Florida Fish and Wildlife Conservation Commission (FWC), Fish and Wildlife Research Institute’s (FWRI) For-Hire At-Sea Observer Survey (here-after, the Observer Program). Observer Program biologists fit discarded gag with Hallprint dart tags, each consisting of a monofilament streamer with a unique identification number, a toll-free reporting hotline, an email contact, and the word “reward”. At the time of release, fish midline length (mm), fishing depth (measured using the vessel’s depth sounder), date and time, geographic location, and hook position (lip, mouth, eye, gill, throat, or gut) were recorded. The program was highly publicized statewide, and anglers returning tags received a free t-shirt to incentivize participation (Sauls 2014). Additional details are provided in Sauls (2014).To align with the spatial extent of the acoustic telemetry experiment and control for potential regional variability in the reporting rate, we limited our consideration to gag released offshore of Tampa Bay, USA (i.e., approximately 85% of all gag analyzed in Sauls 2014, Fig. 1). We converted midline length to total length (TL, mm) using the linear equation provided in Chih (2006) and report this measure to remain consistent with the units used in the acoustic telemetry experiment. All length values hereafter refer to TL unless otherwise specified.

**Figure 1:**
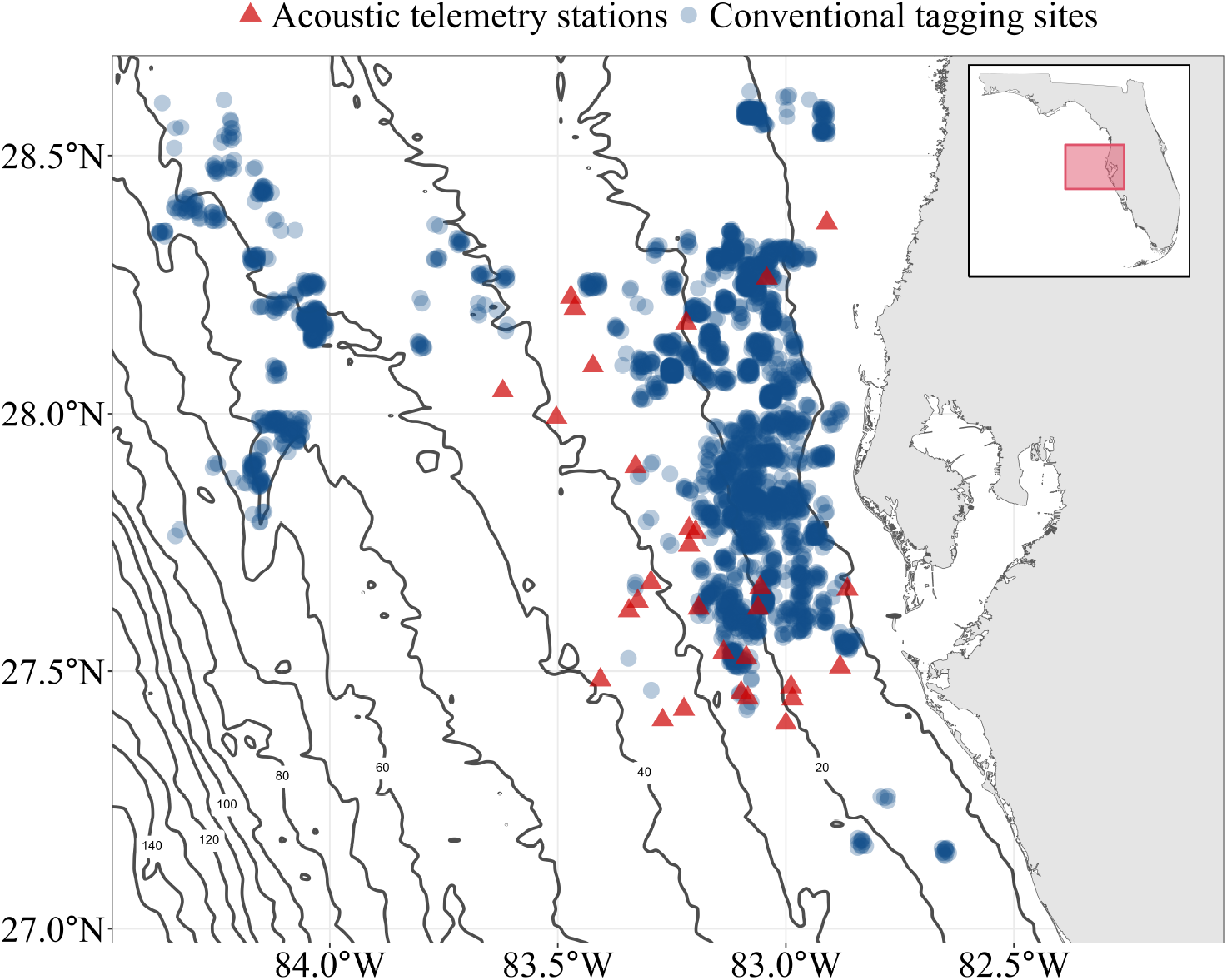
Map of conventionally tagged gag release locations from 2009 through 2012 (blue circles) and acoustic telemetry stations monitored from 2013 through 2017 (red triangles) offshore of Tampa Bay. Darker colors correspond to higher densities of conventional tag releases. Lines show isobaths from 20 m to 140 m at 20 m intervals.

Between December 2013 and February 2017, 90 gag were captured, tagged, and released at acoustic monitoring stations. There were 30 acoustic monitoring stations in total; located 10-90 km offshore of Tampa Bay (5-40 m depth; Fig. 1). Each station contained at least one Vemco VR2W receiver positioned 25-100 m from the station center, with an estimated detection radius of 150-750 m; receivers were serviced and data was downloaded regularly. Fish were captured by experienced recreational offshore anglers instructed to fish as normal. Hook position, TL, and barotrauma symptoms were recorded upon capture. Individuals were fitted with either a V9 transmitter (TL *<* 600 mm; Vemco V9P-2x, 69 kHz) or a V13 transmitter (TL ≥600 mm; Vemco V13P-1x, 69 kHz), along with a Hallprint dart tag, with both tags attached on the same side of the body. Transmitters included pressure sensors that recorded depth at 30-180 s intervals, enabling confirmation of post-release movement and estimation of fish depth when tagged individuals were within receiver range. Fish were either surface-released without venting (N = 23), vented and released at surface (N = 35), or descended with a recompression device (N = 32). To remain consistent with the conventional tagging project, only surface-released fish (i.e., vented or unvented) were considered (N = 58).

Fish that failed to produce detections beyond two days post-release and were not subsequently recaptured were classified as having died post-release. This criterion was selected based on (1) high gag site fidelity (Lindberg et al. 2006), (2) post-release detection patterns of surviving individuals, and (3) prior telemetry studies indicating that acute discard mortality typically occurs within this window for other demersal reef species (e.g., Rudershausen et al. 2025; Zimmermann et al. 2025). In our dataset, the next shortest duration of post-release detection for a surviving fish was nine days, providing additional empirical support that the two-day threshold reliably captured acute mortality without misclassifying survivors (i.e., results were identical across thresholds ranging from two to nine days).

### Analysis

#### Model structure

We modeled post-release survival of both conventionally tagged and acoustically tagged gag via a modified Cormack-Jolly-Seber (CJS) framework (Hyman et al. 2026b). The probability of a tagged fish being recaptured at time *t* is partitioned into: (1) immediate post-release survival (*ψ*); (2) remaining at risk at subsequent time steps (*ϕ*); (3) encounter probability by anglers (*η*); and (4) probability of reporting (*ξ*; Fig 2).

**Figure 2:**
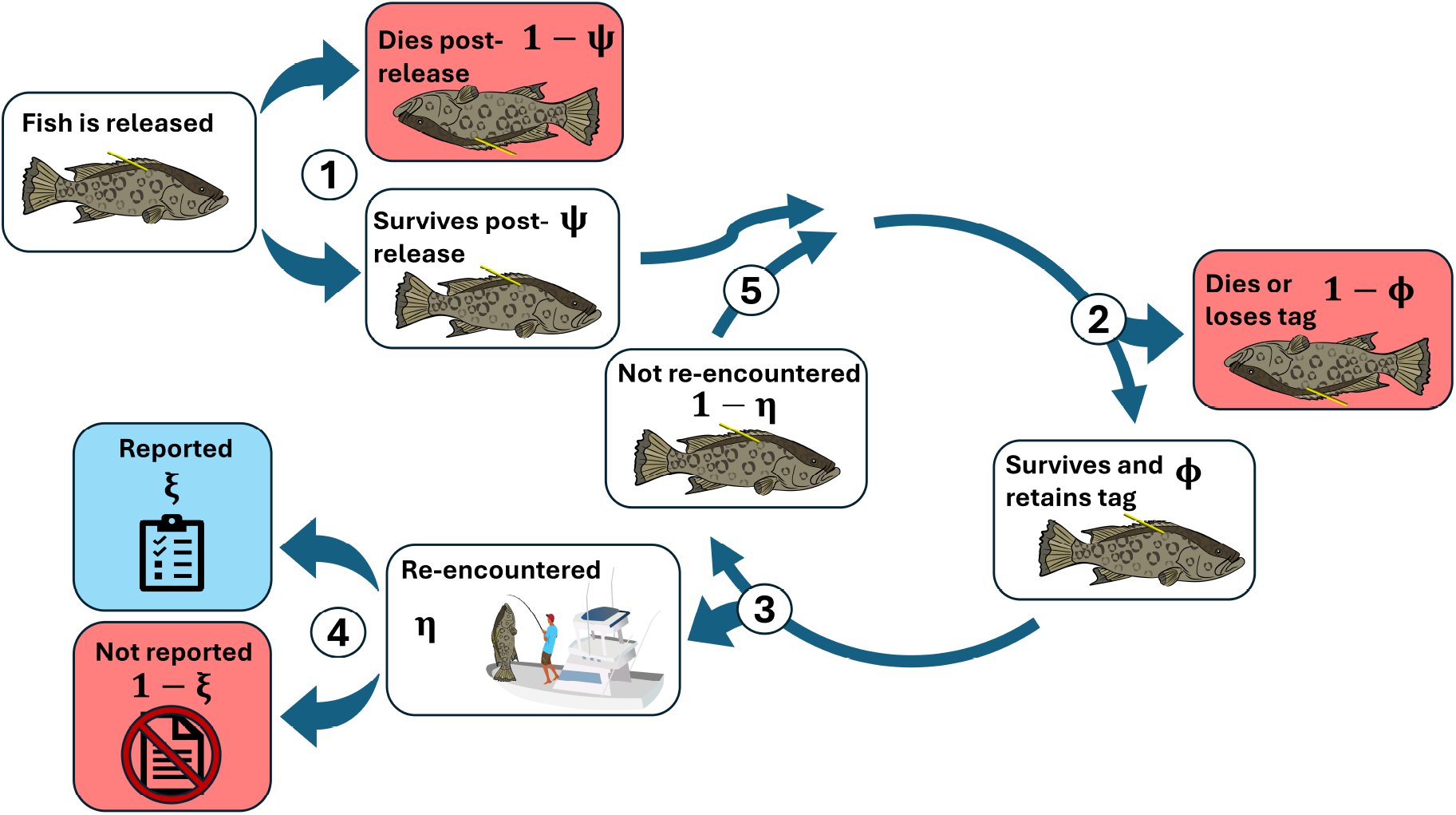
Conceptual diagram modified from (Hyman et al. 2026b) depicting processes determining the fate of a conventionally tagged gag at each time step, resulting in either recapture (*R* = *t*, blue box) or loss (*R* = 0, red boxes). The sequence of events is: (1) survival of the immediate post-release period with probability *ψ*; (2) remain in the risk pool with probability *ϕ*; (3) encountered with probability *η*; (4) if encountered, reported with constant probability *ξ*; or (5) if not encountered, repeat steps 2–5 until encountered, lost, or censored at the end of the study. Graphics were obtained, in part, through the [Integrated Application Network (IAN) Image Library](https://ian.umces.edu/media-library/), and J. Van Vaerenbergh.

In general, for any *t* ≥ *t*_0_, the probability of recapture is:

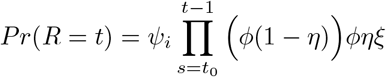

This formulation represents the probability that a fish survives initial release, remains alive and not encountered through time *t* −1, survives natural mortality at time *t*, is encountered by anglers, and is reported. Meanwhile, the probability of never being recaptured (censoring) is the complement of the summed recapture probabilities:

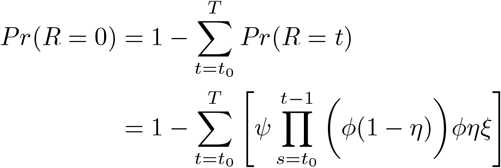

The parameters *ψ, ϕ, η*, and *ξ* can subsequently be estimated from data either as constants or functions of predictors.

We considered the depth of fishing as the primary determinant of post-release survival. Because gag are demersal (e.g., Lindberg et al. 2006; Biesinger et al. 2013), fishing depth was assumed to approximate capture depth; we therefore use “capture depth” hereafter (denoted “Depth” in equations). Although additional variables such as fish length, water temperature, and hook trauma were initially considered as potential predictors, exploratory analyses (Appendix A; Fig S1) indicated that these factors did not explain meaningful variation in post-release survival (acoustic telemetry) or recapture rates (conventional tagging), consistent with previous work (Hyman et al. 2026c). Consequently, these variables were excluded from analysis. Moreover, although venting likely affects post-release survival, angler decisions to vent are usually determined by depth: some anglers vent every fish of a given species once beyond a certain depth, while others vent only when external barotrauma symptoms are present, which are more prominent in deeper waters. Indeed, in both experiments, venting was positively correlated with depth. Because our objective was to estimate the total effect of depth (both direct and indirect) rather than isolate venting effects, including this variable would induce post-stratification bias.

For both data types, post-release survival, *ψ*, was modeled as a function of capture depth using a logit link to constrain values to the probability scale:

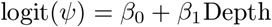

Here, *β*_0_ denotes baseline survival in logit space, and *β*_1_ represents the effect of capture depth on post-release survival. We assumed baseline post-release survival followed a Beta(15, 1) prior by adopting a non-informative uniform prior – i.e., Beta(1, 1) – and updating it using directly observed post-release survival trials from previous work(Burns et al. 2002). This distribution was approximated with a normal distribution in logit space centered at 3.25 with a standard deviation of 1.31. For *β*_1_, logistic regression results from 3,876 conventionally tagged gag released along the Atlantic coast of the United States estimated a depth effect of −0.032 (SE = 0.004; McGovern et al. 2005). We assigned a prior of *N* (− 0.032, 0.04), inflating the reported standard error by a factor of ten to reflect uncertainty in transferability across regions and study designs. Sensitivity analyses evaluating weaker priors (i.e., variance = 4) for both beta coefficients did not appreciably change our results, and we therefore concluded that the model was robust to modest variation in the choice of priors.

For acoustic telemetry data, post-release survival for each fish *j* was represented as a binary variable (*y*_*j*_; where 1 = survival and 0 = death), and modeled explicitly using logistic regression:

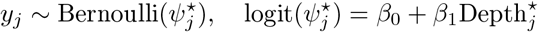

The parameter vector 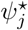 is meant to differentiate post-release survival probabilities estimated for the *j*^*th*^ acoustically tagged fish from estimates corresponding to *i*^*th*^ conventionally tagged fish (i.e., *ψ*_*i*_).

Notably, because the post-release survival probabilities of conventionally tagged fish *ψ*_*i*_ and the reporting rate *ξ* enter the likelihood as a product, these parameters are structurally non-identifiable without direct fate observations of post-release survival provided by acoustically tagged fish. By parameterizing post-release survival with shared *β* coefficients across both tag types, the reporting rate *ξ* becomes separately estimable on the logit scale. The parameter *λ* – the logit-transformed reporting rate – was assigned a *N* (0, 1.5) prior, corresponding to a flat prior in 0-1 probability space.

For conventionally tagged fish, the probability of remaining at risk of recapture (i.e., surviving with an intact tag; *ϕ*) was modeled via the relationship *ϕ* = exp(− *M*), where *M* denotes instantaneous monthly natural mortality (e.g., Williams-Grove and Szedlmayer 2016). Literature-derived mortality estimates were incorporated by placing a prior on the parameter *γ* corresponding to log(*M*) using a log-log link. The most recent stock assessment estimated annual instantaneous natural mortality for Gulf gag aged approximately 2-7 years (consistent with the size range of conventionally tagged individuals) between 0.19 and 0.36 (SEDAR72 2021). Assuming constant mortality throughout the year, these values were converted to monthly rates. The log-transformed monthly mortality estimates yielded a mean of −3.88 with a standard deviation of 0.23, which was used as the prior for *γ*.

To separate variation in encounter rates (*η*) from true post-release survival in our conventional tagging data, we divided the study area into 0.25^◦^ *×* 0.25^◦^ latitude–longitude cells (≈ 682 ^2^ km each), yielding *L* = 20 spatially distinct locations. Time was discretized into monthly intervals (*T* = 44). Fish at large beyond December 2012 were censored. Seasonal variation in effort was modeled using harmonic regression terms (i.e., sine- and cosine-transformed temporal data denoted *x*) for annual and semi-annual periodicities (Trudeau et al. 2022; Hyman et al. 2024; Hyman et al. 2026a). Spatial heterogeneity in effort was captured via location-specific random effects *τ*_*l*[*i*]_. The harmonic terms allow the model to capture predictable seasonal variation in recreational fishing, while random effects account for consistently higher or lower activity in particular areas (e.g., near urban centers vs offshore). The linear predictor was constrained to the 0-1 probability scale using a complementary log-log link. The hazard of encounter for a given fish is scaled by size-selectivity *σ*_*i*_ to reflect size-dependent vulnerability to recapture. Aggregate selectivity was estimated based on recreational selectivity curves for circle hooks as a function of gape-size, weighted to the distribution of hook sizes recorded by the Observer Program during the time-frame and location of the conventional tagging study (details provided in Appendix B; Fig S2).

The full model for the probability of recapture for conventionally tagged fish *i* at time *t* in location *l*[*i*], and acoustically tagged fish *j* is therefore:

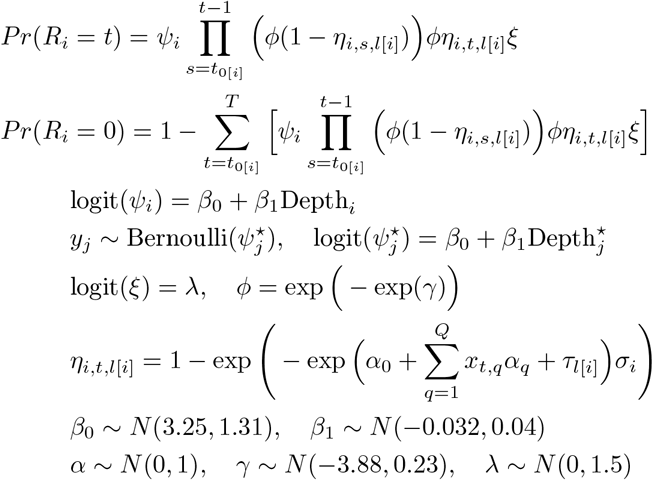

#### Model implementation and validation

All models were implemented in Stan (Gelman et al. 2015; Stan Development Team 2022) via R. Four chains were run with 1,000 warm-up and 1,000 sampling iterations each, yielding 4,000 posterior draws. Convergence was assessed via 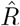 (≈ 1 at convergence). Covariates were considered meaningful if 80% credible intervals (CIs) excluded zero. Unless otherwise noted, reported values are maximum *a posteriori* (MAP) estimates with 80% CIs. Simulation-based validation was conducted to evaluate the model’s ability to recover known parameter values under conditions reflecting the structure and sampling characteristics of the observed data (Appendix C). Posterior predictive checks were subsequently performed to assess model performance by (1) comparing in-sample predictions with observed recapture frequencies and post-release survival outcomes and (2) comparing estimated probabilities of encounter to local estimates of angler effort for reef fish (derived from Hyman et al. 2024, Appendix C).

#### Projections

Using Observer Program data as a proxy for the recreational fleet and posterior draws from our model, we estimated monthly and annual post-release survival during 2023, 2024, and 2025. For each recorded gag discard *p*, the depth of capture Depth_*p*_ and posterior draw *d* were used to compute the post-release survival probability 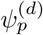 as:

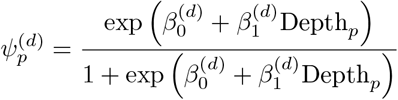

We estimated the average post-release survival for each month *m* and posterior draw *d* by averaging predicted probabilities across all observations *p* within that month:

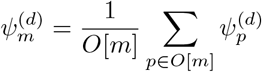

where *O*[*m*] denotes the set of observations in month *m*. Repeating this calculation across all posterior draws yields the full posterior distribution of monthly post-release survival probabilities, *ψ*_*m*_. We repeated this procedure at the annual resolution for each year to obtain the full posterior distribution of post-release survival for years 2023, 2024, and 2025.

## Results

For conventionally tagged gag, capture depth ranged from 3 to 93 m (median = 19 m), and total length ranged from 248 to 1012 mm (median = 490 mm). Acoustically tagged gag were captured at depths between 6 and 37 m, with total lengths ranging from 443 to 803 mm. Among conventionally tagged individuals, 356 fish were recaptured and reported, corresponding to a 10.2% return rate. The median time at large for recaptured fish was 57 days (range: 1–1,058 days). Of the 58 acoustically tagged fish considered, 54 survived post-release (93.1% survival). Both recapture proportions (conventional tags) and post-release survival (acoustic tags) declined with increasing capture depth (Fig. 3). No post-release mortality was observed among acoustically tagged gag released at depths shallower than 25 m.

**Figure 3:**
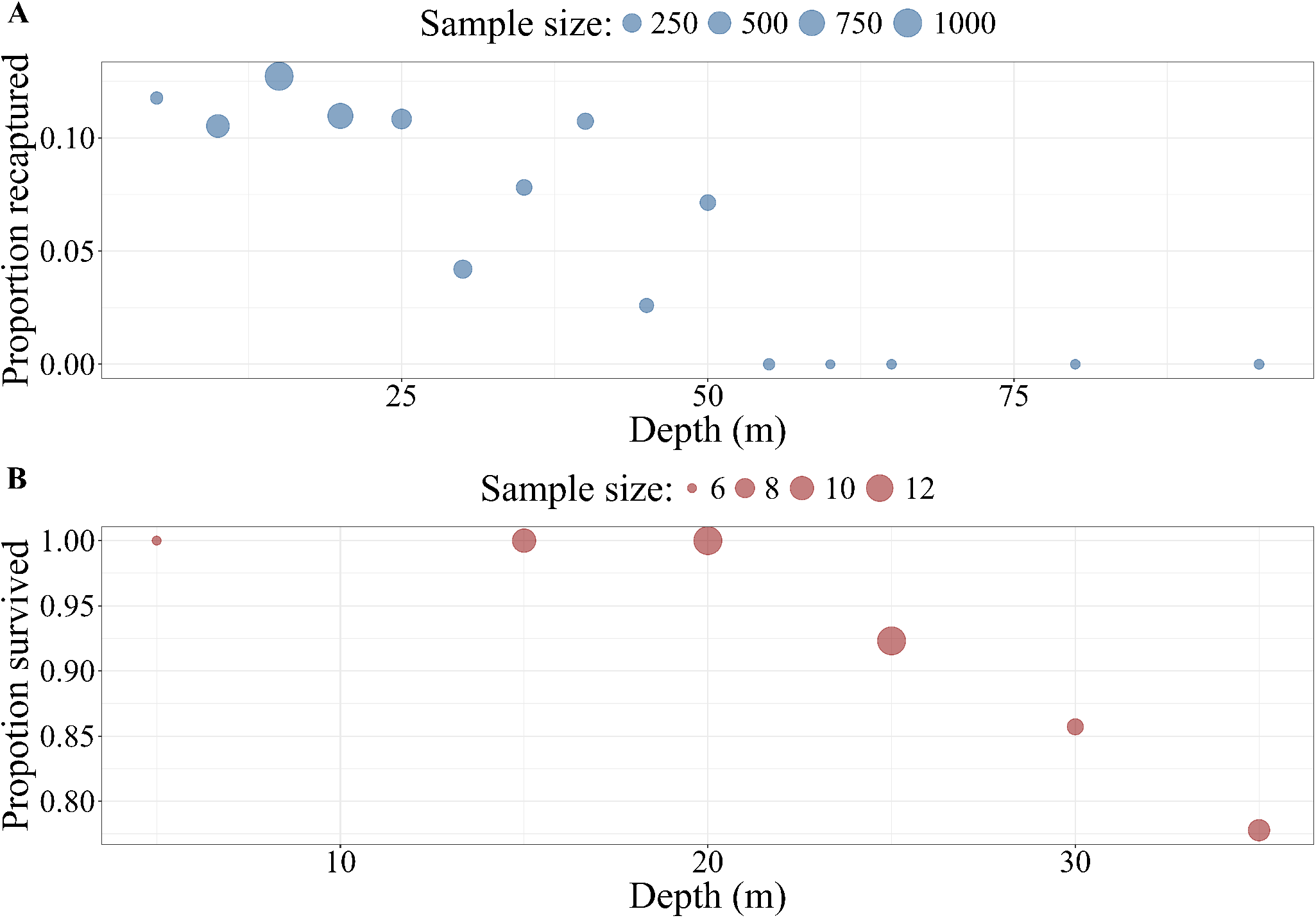
Proportions of gag that A) were recaptured or B) survived post-release as a function of depth (binned to 5 m intervals) based on raw conventional tagging and acoustic telemetry data, respectively. Point size is relative to the number of fish in each depth bin, which varies based on experiment.

Validation and diagnostic exercises indicated model inferences were robust. Simulations indicated that the model framework reliably captured true parameter estimates in posterior distributions (Fig. S3), and posterior predictive checks showed strong agreement with the observed data (Fig. S4).

Moreover, estimated seasonal variation in encounter probability was consistent with independent estimates of local recreational fishing effort, providing additional support that processes pertaining to post-release survival, encounter, and reporting were separable (Fig. S5).

Model results supported the depth-related patterns observed in the raw data. The MAP estimate of *β*_0_ corresponded to a baseline post-release survival probability of 0.97 (80% CI: 0.94–0.98). The MAP estimate of *β*_1_ was negative, and its 80% CI excluded zero, indicating that post-release survival decreased meaningfully with increasing capture depth (Table 1; Fig. 4). At 93 m, the maximum capture depth observed, predicted post-release survival declined to 0.32 (80% CI: 0.12–0.64). The MAP estimate of *ξ* (reporting rate) and *ϕ* (probability of remaining at risk) were 0.14 (80% CI: 0.13–0.15) and 0.98 (80% CI: 0.97-0.99), respectively.

**Table 1:** Posterior summary statistics – maximum *a posteriori* (MAP) and 80% credible interval – of the predictor coefficients of the integrated tagging model. All values are rounded to two decimal places.

| Predictor | 10% | MAP | 90% |
| --- | --- | --- | --- |
| $\beta_0$ | 2.76 | 3.36 | 3.95 |
| $\beta_1$ | -0.06 | -0.05 | -0.03 |
| $\xi$ | 0.13 | 0.14 | 0.15 |
| $\phi$ | 0.97 | 0.98 | 0.99 |
| $\alpha_0$ | -2.25 | -1.90 | -1.69 |
| $\alpha_1$ | -0.10 | 0.01 | 0.14 |
| $\alpha_2$ | -0.05 | 0.04 | 0.15 |
| $\alpha_3$ | -0.56 | -0.44 | -0.35 |
| $\alpha_4$ | 0.19 | 0.31 | 0.40 |

**Figure 4:**
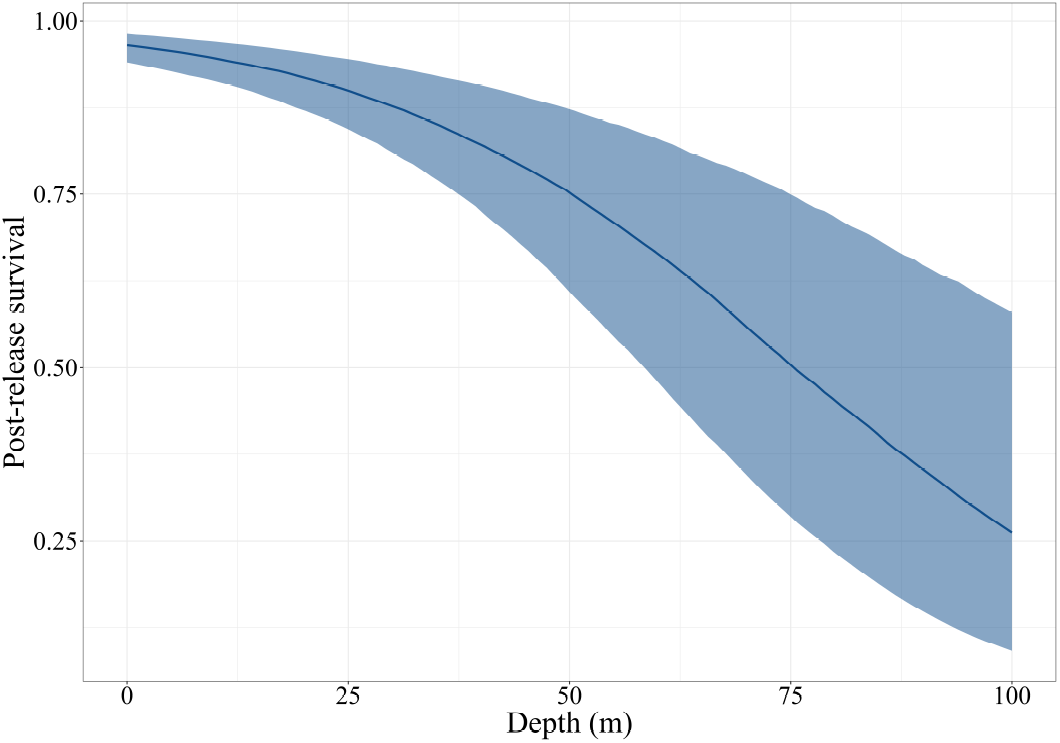
Predicted gag post-release survival rates as a function of capture depth. The solid line denotes the maximum *a posteriori* post-release survival estimate, whereas the band denotes the 80% credible interval. Higher uncertainty at deeper capture depths corresponds with smaller sample sizes in conventional tagging data.

Monthly post-release survival rates, predicted from capture depth, exhibited clear seasonal variation corresponding to changes in angler fishing behavior. Based on Observer Program data (2023–2025), mean capture depths for discarded gag were lowest in fall and winter (September-January); typically between 10 and 20 m (Fig. 5). These depths corresponded to MAP post-release survival estimates of approximately 0.93. Following February, mean capture depth increased, reaching a seasonal maximum of 38 m in July, which corresponded to a MAP survival estimate of 0.83. Annual MAP estimates of post-release survival were 0.86 (80% CI: 0.80–0.93), 0.92 (80% CI: 0.88–0.95), and 0.87 (80% CI: 0.81–0.93) for 2023, 2024, and 2025, respectively.

**Figure 5:**
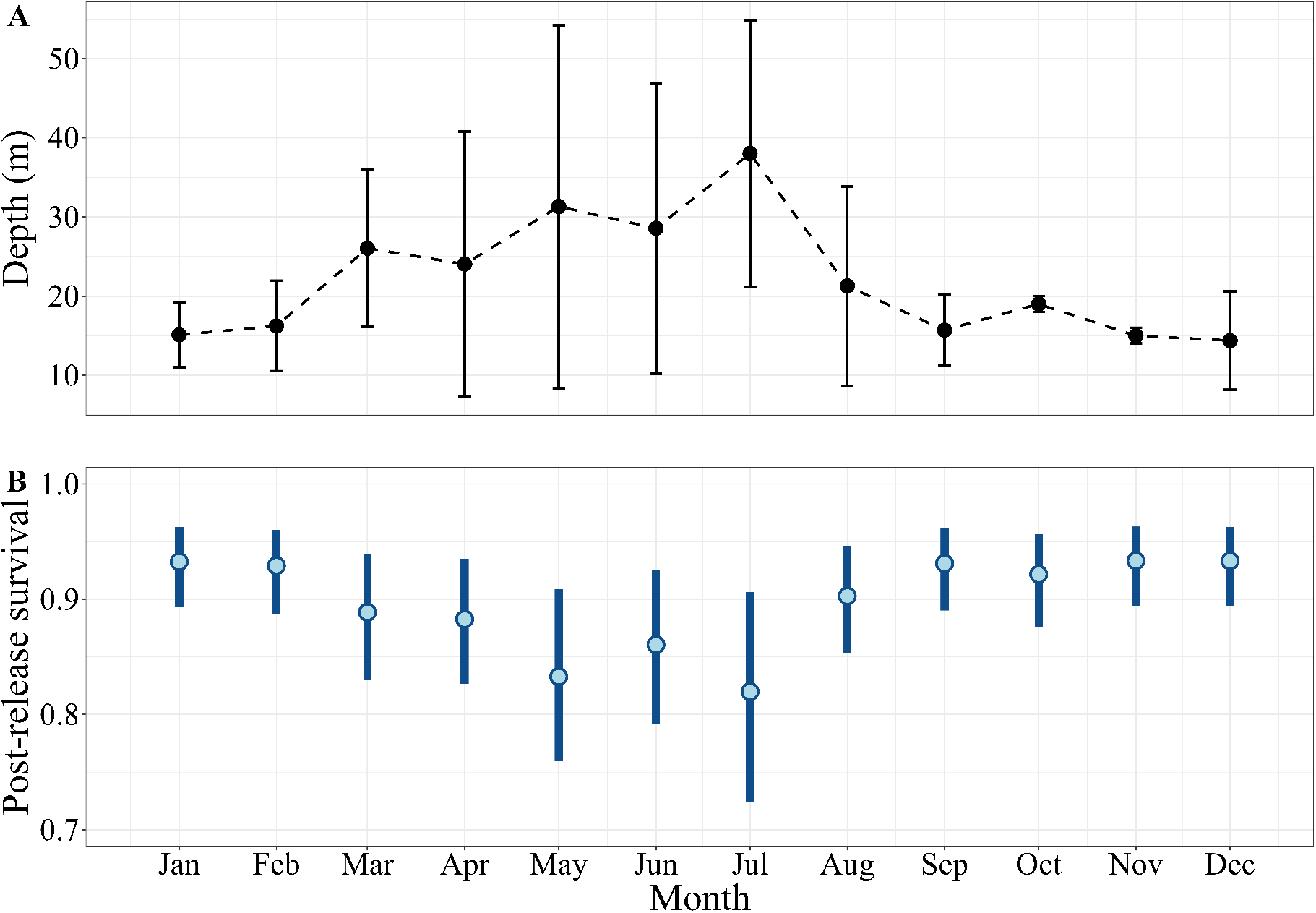
Seasonal variation in gag post-release survival probability estimated from Observer Program data (2023–2025). (A) Median monthly capture depth of released gag, with error bars representing standard error. (B) Posterior predictive distributions of monthly post-release survival based on observed capture depths. Circles denote the maximum *a posteriori* predictive estimate, while bars denote the 80% credible interval.

## Discussion

Integrating telemetry-derived fate information with large-scale conventional tagging data within a unified statistical framework addresses key limitations inherent to each methodology when applied independently (Benoît et al. 2020). Our approach enabled the estimation of post-release survival across a broad range of environmental conditions and as a function of the specific conditions under which recreational fishing occurs. Model results, applied to observed depth of capture and release from the recreational for-hire fleet, produced estimates that are broadly consistent with values assumed in recent stock assessments and indicate high survivorship of recreationally released gag. More broadly, this framework provides a tractable approach for estimating post-release survival in highly targeted fisheries, particularly those where regulatory discards frequently exceed retained catch.

Combining acoustic telemetry with conventional tagging improved inference on gag post-release survival by addressing the limitations associated with each method when used alone. For example, Sauls (2014) analyzed the same conventional tagging dataset to estimate relative survival of gag across a wide depth range but could not distinguish discard mortality from imperfect reporting, instead providing absolute survival estimates based on assumed baseline post-release survival rates of 100%, 92.5%, and 85%. Conversely, absolute post-release survival could be estimated from the acoustic telemetry dataset alone, but only over a limited geographic region and range of capture depths. By applying a statistical model to a subset of the tagging data in combination with acoustic telemetry observations from the same area and general time frame, our integrated approach enabled the estimation of post-release survival across the full range of environmental conditions encountered by the fishery while grounding baseline values in directly observed outcomes.

Patterns of post-release survival estimated in this study are consistent with evidence from cage experiments and post-release condition assessments of discarded gag. For example, Burns et al. (2002) reported 100% survivorship of gag captured at 30 m and monitored in holding cages, indicating a high capacity for survival following capture from moderate depths under controlled conditions. In addition, observer data from 1,650 gag discarded by anglers aboard for-hire vessels between 2022 and 2024 indicated that 81% were released without major external signs of barotrauma and were able to return rapidly to depth (Hyman et al. 2026c), suggesting high post-release survival. These observations align with our acoustic telemetry results, in which no post-release mortality was observed among the 32 gag released at depths ≤ 25 m. Despite high survivorship at shallower depths, post-release survival declined with increasing depth, consistent with previous studies documenting elevated barotrauma incidence and impairment with greater capture depth for gag (Sauls 2014; Hyman et al. 2026c) and demersal species more broadly (e.g., Burns 2009; Ferter et al. 2015; Runde et al. 2019; Bohaboy et al. 2020; Ramsay et al. 2025).

Applying the results of our integrated model to observations from the recreational fishery enabled estimation of post-release survival probabilities averaged across the environmental conditions and fishing practices actually experienced by released gag, providing the absolute post-release survival inputs required for stock assessments. Assuming that Observer Program records are representative of both for-hire and private-recreational fleets, survival probabilities for individual fish, along with associated uncertainty, can be estimated based on the environmental conditions at capture (Hyman et al. 2026b). Averaging these values across all fish produces robust estimates of overall post-release survival, while the bounds of uncertainty can be incorporated into assessments through sensitivity analyses to evaluate how variability in survival affects stock status determinations. Notably, our annually averaged estimates of post-release survival from 2023, 2024, and 2025 (86%, 92% and 87%, respectively) align with the recreational post-release survival rate used in the most recent Gulf gag stock assessment (i.e., 88%; SEDAR72 2021), providing independent support for recent stock status determinations.

In addition to fishery-wide averages, our results reveal temporal patterns with potential management relevance. Monthly estimates indicate lower post-release survival during summer months, corresponding to fishing in deeper water. This information can guide outreach and mitigation efforts; for example, targeting education on descending device use during periods when anglers fish farther offshore may reduce discard mortality. Such insights are particularly important for vulnerable stocks, such as Gulf gag, which is currently under a rebuilding plan following its recent designation as impaired (SEDAR72 2021). Results also support the recent decision to shift the recreational gag season start date from June to September, which likely reduced overall dead discards (Hyman et al. 2025).

More broadly, our integrated approach may be useful for estimating post-release survival of other highly targeted species. When absolute post-release survival rates under specific environmental and fishing conditions are well-estimated using abundant acoustic telemetry data (e.g., red snapper, *Lutjanus campechanus*; Bohaboy et al. 2020; Runde et al. 2021; Rudershausen et al. 2025; Zimmermann et al. 2025), conventional tagging studies can incorporate these estimates as auxiliary information, priors within a Bayesian framework, or as supplied data to anchor relative risk estimates. When such information is unavailable, the simultaneous deployment of conventional and electronic tags within the same geographic region provides the opportunity to robustly estimate relationships between post-release survival and relevant predictors across a broad range of environmental conditions. Finally, monitoring data are critical for applying experimentally derived relationships to the specific conditions under which fishing occurs, thereby refining post-release survival estimates used in stock assessments and revealing spatiotemporal patterns in survivorship that can inform management decisions.

### Limitations and future work

Although our methodology represents an advancement in estimating post-release survival, inherent limitations remain. First, the recapture model framework assumes that all tagged fish encountered by anglers but not reported are removed from the risk pool (i.e., harvested or released without a tag; Hyman et al. 2026b). While this assumption is implicit in all relative-risk frameworks, including logistic and Cox regression, it is unrealistic because some fraction of unreported fish is likely re-released with intact tags. If environmental conditions during subsequent releases are similar to those at initial capture and release, mortality risk may compound. For demersal species such as gag, which exhibit high site fidelity, fish captured in deeper waters during repeated encounters are increasingly susceptible to barotrauma, potentially exaggerating the relationship between depth and post-release survival and underestimating overall survival rates. Although the relatively high post-release survival observed in our study reduces the practical impact of this bias for gag, it could be more consequential for other highly targeted species. Future modeling frameworks that relax this assumption would improve postrelease survival estimation when conventional tag-return data are used.

Second, our estimates rely on the assumption that discard practices recorded by the Observer Program are representative of the broader private-recreational fleet. This assumption is commonly made in the absence of private-recreational data, but estimates may be biased if fishing practices, such as average fishing depth, differ substantially between the for-hire fleet (well-represented in the Observer Program) and the larger private-recreational fleet. Quantifying the degree to which this assumption is valid would improve confidence in post-release survival estimates and strengthen their application to stock assessment and management.

## Acknowledgments

Conventional tagging would not have been possible without the dedication and sustained effort of FWC observers, volunteer for-hire captains and crew, and FWC staff who coordinate and administer the FWC tag-return program. Acoustic telemetry tagging and data collection were made possible through the support of the St. Petersburg Underwater Club, as well as numerous for-hire captains and private recreational anglers who contributed their time and expertise. Methodological refinement was substantially strengthened through the input of B. Sauls. This work represents a contribution from the Quantitative Fisheries Assessment Workgroup within the Center for Environmental Analysis, Synthesis, and Application and reflects a collaborative effort between the University of South Florida and the Florida Fish and Wildlife Research Institute.

## Appendix A: Exploratory data analysis and predictor selection

We employed a combination of model selection and exploratory data analysis (EDA) to determine the final predictors of post-release survival in our integrated discrete-time framework. First, we evaluated predictive performance using acoustic telemetry data only, applying the estimated log pointwise predictive density (ELPD) and corresponding ΔELPD values, which quantified the difference in predictive accuracy between each candidate model and the best-performing model (Vehtari et al. 2017). ELPD reflects out-of-sample predictive accuracy and was estimated via the approximate Leave-One-Out Information Criterion (LOO-IC) (Gelman et al. 2015; Vehtari et al. 2017) using the loo package (Vehtari et al. 2022). When ΔELPD values were small (≤ 4), we selected the simpler model for parsimony (Sivula et al. 2020).

Immediate post-release survival (1 = survived, 0 = died) was initially modeled using logistic regression through the brms package (Bürkner 2017). Candidate predictors included capture depth, hook trauma (presence/absence of hooks in eyes, throat, gills, or gut), total length, and the absolute difference between surface and bottom water temperatures at the time of capture (Table S1).

Model selection indicated that depth alone (*g*_0_) was the most parsimonious predictor of post-release survival. The MAP estimate of the depth coefficient was negative (−0.20) and its 80% credible interval excluded zero (−0.33 to −0.09), confirming a meaningful negative relationship between capture depth and post-release survival.

EDA of conventional tagging data supported this conclusion (Fig. S1). Scatter plots of recapture rates versus water temperature differences revealed no discernible pattern. While recapture probability declined noticeably with depth, rates among fish with and without hook trauma were similar, and apparent size-related patterns in recapture rates were consistent with the recreational selectivity curve (Appendix B), rather than reflecting true survival differences. These results corroborate prior work (Hyman et al. 2026c), which found no detectable effects of length, hook trauma, or temperature on gag release condition, but identified depth as a key driver of post-release outcomes for this species.

Although the acoustic telemetry dataset was modest (N = 58), posterior distributions were stable, and patterns aligned with trends observed in the larger conventional tagging dataset. We therefore concluded that capture depth was the primary determinant of post-release survival in gag, and selected it as the sole predictor in the integrated discrete-time model.

## Appendix B: Selectivity estimation

The probability of encountering a tagged fish is influenced by gear selectivity, whereby individuals of certain sizes may be less likely to be captured, all else being equal, for a given hook size (e.g., Campbell et al. 2014; Christiansen et al. 2020; Christiansen et al. 2022). Conditioned on fishing effort and remaining at risk, the relative probability that a fish of length *L* is encountered, compared with the length at which encounter probability is maximized, is defined as:

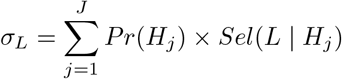

where *Pr*(*H*_*j*_) denotes the probability that an angler uses hook size *H*_*j*_, and *Sel*(*L* | *H*_*j*_) represents the relative selectivity of hook size *H*_*j*_ for fish of length *L*.

To account for size-related bias in encounter probability, we incorporated selectivity estimates derived from the distribution of hook sizes used in the recreational fishery. Specifically, we constructed an aggregate selectivity curve as a proxy for *σ*_*L*_ based on: (1) a lognormal selectivity function describing *Sel*(*L*_*i*_ | *H*_*j*_) as a function of hook gape size, and (2) the proportion of fishing effort associated with each hook gape size, estimated from Observer Program data collected off Tampa Bay between 2009 and 2012, used here as a proxy for *Pr*(*H*_*j*_).

The lognormal selectivity function followed the formulation of Christiansen et al. (2020), who estimated gear selectivity for red grouper (the closest available morphometric analogue to gag) on the west coast of the Florida Peninsula using fishery-independent survey data (Fig. S2A). Aggregate relative selectivity *σ*_*L*_ was then calculated as the weighted sum of hook-specific selectivity curves, with weights corresponding to the observed proportion of effort allocated to each hook gape size (Fig. S2B). The resulting curve was normalized to a maximum value of 1 (Fig. S2C) and used to scale the hazard of encounter within the modeling framework.

## Appendix C: Model validation and diagnostics

Prior to formal analysis, we conducted a simulation study to evaluate the model’s ability to recover known parameter values under conditions reflecting the structure and sampling characteristics of the observed datasets. Using the empirical distributions of release conditions, locations, and times for both conventionally and acoustically tagged gag, we generated a synthetic dataset consistent with the proposed data-generating process, with parameter values fixed at known quantities.

To ensure that parameter recovery was not artificially facilitated by prior specification, simulated data were fit using a version of the model with weakly informative priors (i.e., *N* (0, 2) for all estimable parameters). Across simulations, posterior estimates for fixed-effects parameters were generally centered near their true values, and the true parameter values were consistently contained within all but one of the corresponding 80% credible intervals and all 95% intervals (Fig. S3). These results indicate that the model is capable of reliable parameter recovery and does not exhibit evidence of structural bias under realistic sampling conditions.

Following model fitting to the observed data, posterior predictive checks (PPCs) were conducted to evaluate whether model predictions were consistent with observed values. For each conventionally tagged gag and for each combination of posterior draws, we simulated a time-to-recapture value. We assessed model performance by comparing the overall proportion of fish recaptured, examining predicted and observed recapture probabilities grouped into evenly spaced bins, and comparing the empirical distribution of observed recapture times with distributions generated from 1,000 posterior predictive draws. In addition, the empirical cumulative distribution function (ECDF) of observed recapture times was compared with ECDFs simulated from posterior predictive draws (Fig. S4). Across all diagnostics, the results indicated a high degree of agreement between observed recaptures and model predictions.

In addition to PPCs, we evaluated seasonal encounter probabilities based on the posterior distributions of the *α* parameters against independent estimates of local recreational reef-fishing effort (Hyman et al. 2024), which are expected to correlate with encounter probabilities (e.g., Hyman et al. 2026b). For simplicity, spatial random effects were fixed at zero. Posterior predictive distributions of seasonal encounter probabilities showed peaks in spring and fall, consistent with observed patterns in angler trips (Fig. S5), providing additional support that the model successfully partitioned post-release survival and encounter probabilities.

## Appendix D: Supplemental tables and figures

**Table S1:** Selection results for logistic regression models (*g*_*m*_) predicting gag post-release survival from acoustic telemetry data, ranked by predictive performance using the approximate Leave-One-Out information criterion (**LOO**). Reported values include **ELPD**_**LOO**_, the estimated log pointwise predictive density, and **Δ**_**ELPD**_, the difference between each model and the best-performing model. Covariates include capture depth (**Depth**), hook trauma (**Hook**), total length (**Length**, mm), and the absolute surface–bottom temperature difference at capture (**ΔTemp**). The selected model (*g*_0_) is shown in bold.

| Model | Formula | LOO | $ELPD_{LOO}$ | $\Delta_{ELPD}$ |
| --- | --- | --- | --- | --- |
| $g_1$ | Depth + Hook | 24.25 | -12.13 | 0.00 |
| $g_2$ | Depth + Hook + Length | 25.53 | -12.76 | -0.64 |
| **$g_0$** | **Depth** | **26.45** | **-13.22** | **-1.10** |
| $g_3$ | Depth + Hook + Length + $\Delta\text{Temp}$ | 29.23 | -14.62 | -2.49 |

**Figure S1:**
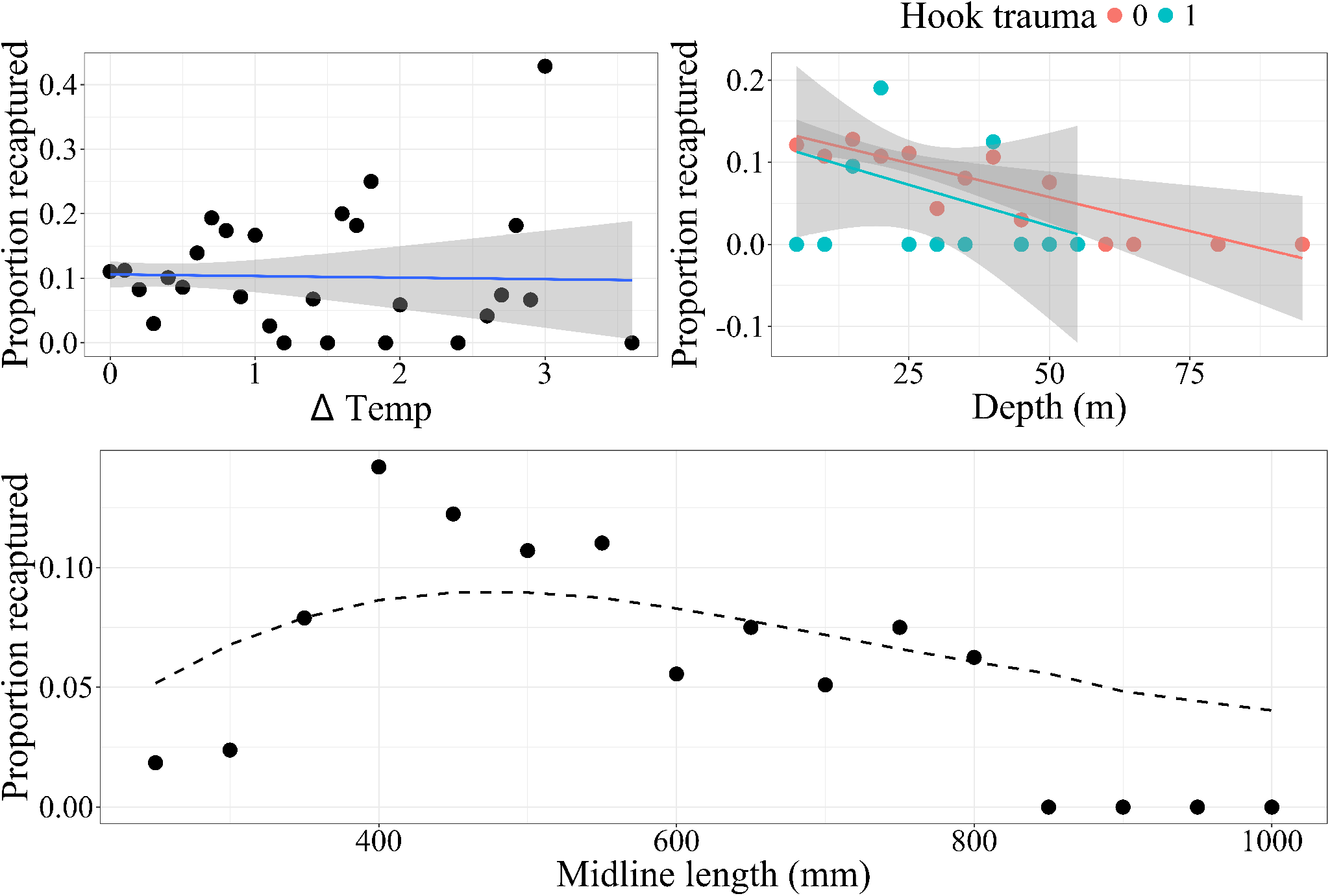
Exploratory data analysis of recapture rates in the conventional tag dataset. Top-left: difference between surface and bottom-water temperatures; top-right: depth (5 m bins) grouped by presence (1) or absence (0) of hook trauma; bottom: fish size (50 mm bins), with the applied selectivity curve (Appendix B) scaled and superimposed. Linear regression lines are overlaid where appropriate.

**Figure S2:**
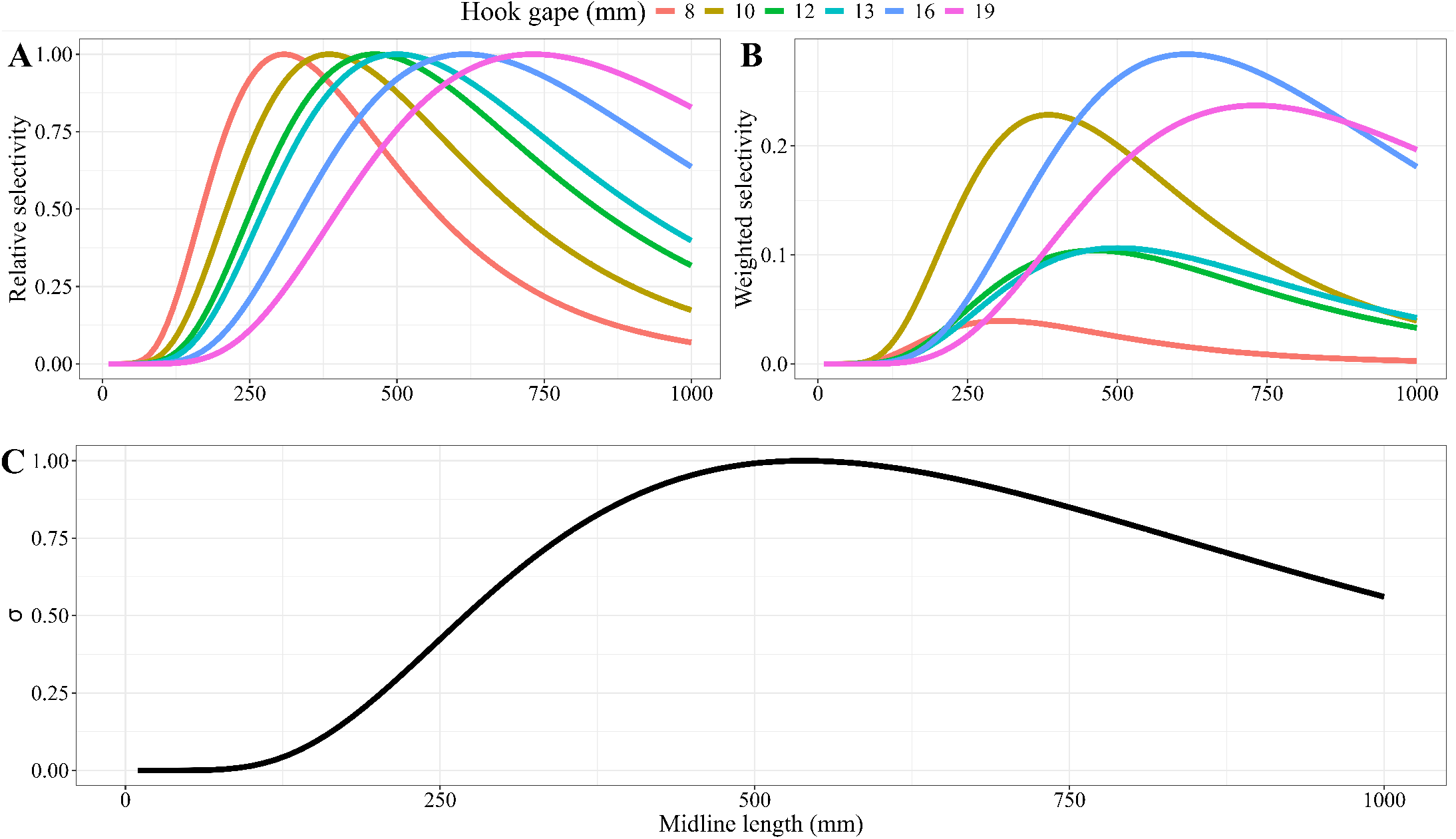
Plots depicting aggregate relative size-selectivity curve (*σ*) estimation: A) Relative selectivity curves from each hook gape size recorded in the Observer Program between 2009 and 2012, filtered for reef fish trips of Tampa Bay, based on model outputs from Christiansen et al. (2020); B) the same relative selectivity curves with peaks scaled based on the proportion of total effort spent fishing with each hook size; C) the resulting aggregate selectivity curve based on the weighted sum of all six relative selectivity curves and standardized to have a maximum value of one. Values refer to midline (fork) length to remain consistent with Christiansen et al. (2020).

**Figure S3:**
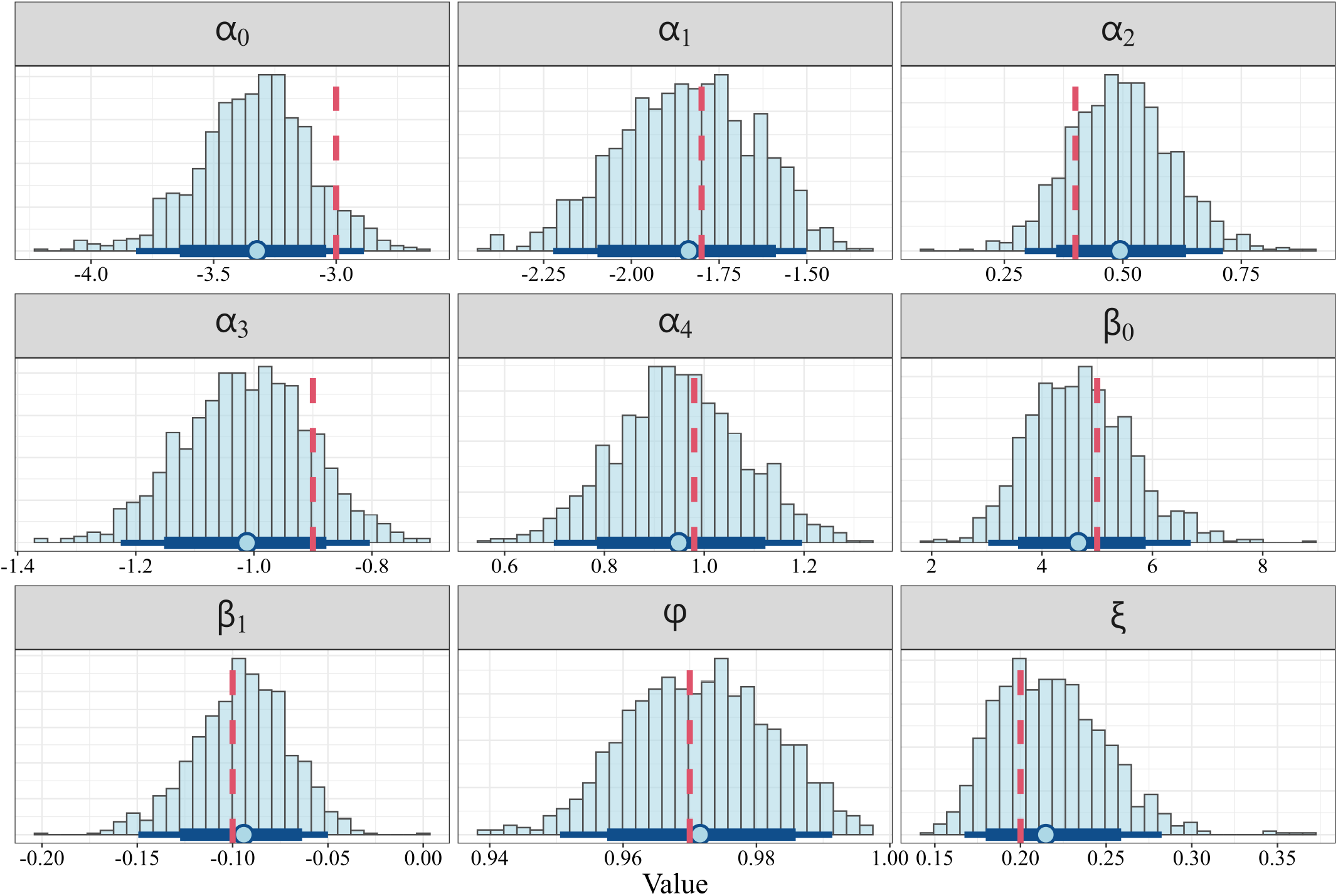
Posterior distributions of fixed-effect parameters obtained from the simulation-based parameter recovery analysis. Histograms represent full posterior distributions. Points indicate maximum *a posteriori* estimates, thick dark blue bands denote the corresponding 80% credible intervals, and thin dark bands correspond to 95% credible intervals. Red dashed lines mark the true parameter values used to generate the simulated data.

**Figure S4:**
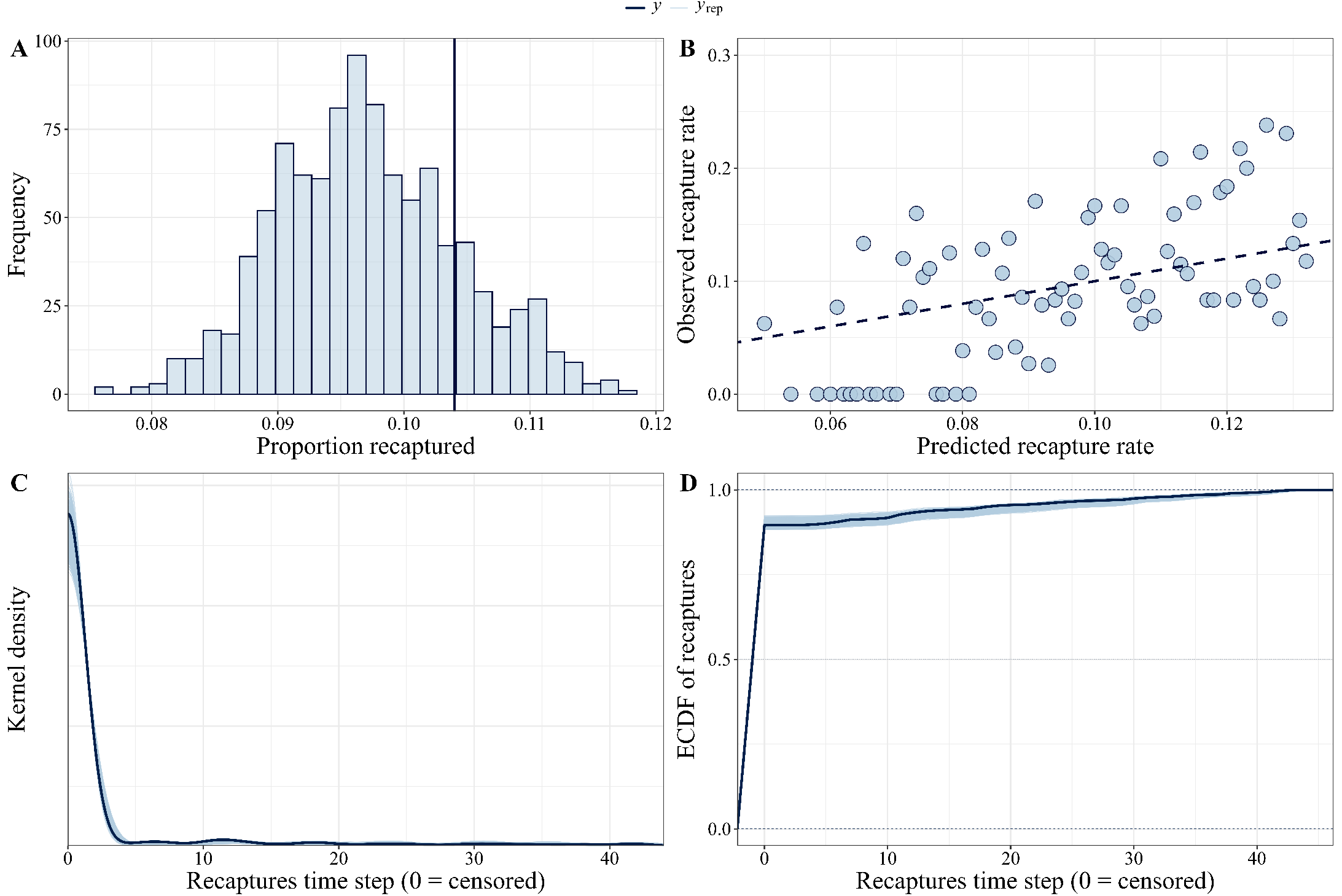
Posterior predictive checks based on conventional tagging data. (A) Distribution of simulated recapture proportions with the observed recapture proportion indicated by the vertical line. (B) Posterior predictive calibration plot showing agreement between predicted and observed recapture probabilities. Predicted probabilities were grouped into evenly spaced bins, with the mean posterior predicted probability (x-axis) compared against the observed proportion of recaptures (y-axis) in each bin. The dashed 1:1 line denotes perfect calibration; deviations reflect systematic bias in predicted recapture probabilities. (C) A comparison of the empirical distribution of observed recapture times to the distributions of 1,000 scans from the posterior predictive distribution. (D) Empirical cumulative distribution function (ECDF) of observed recapture times compared with ECDFs simulated from posterior predictive draws.

**Figure S5:**
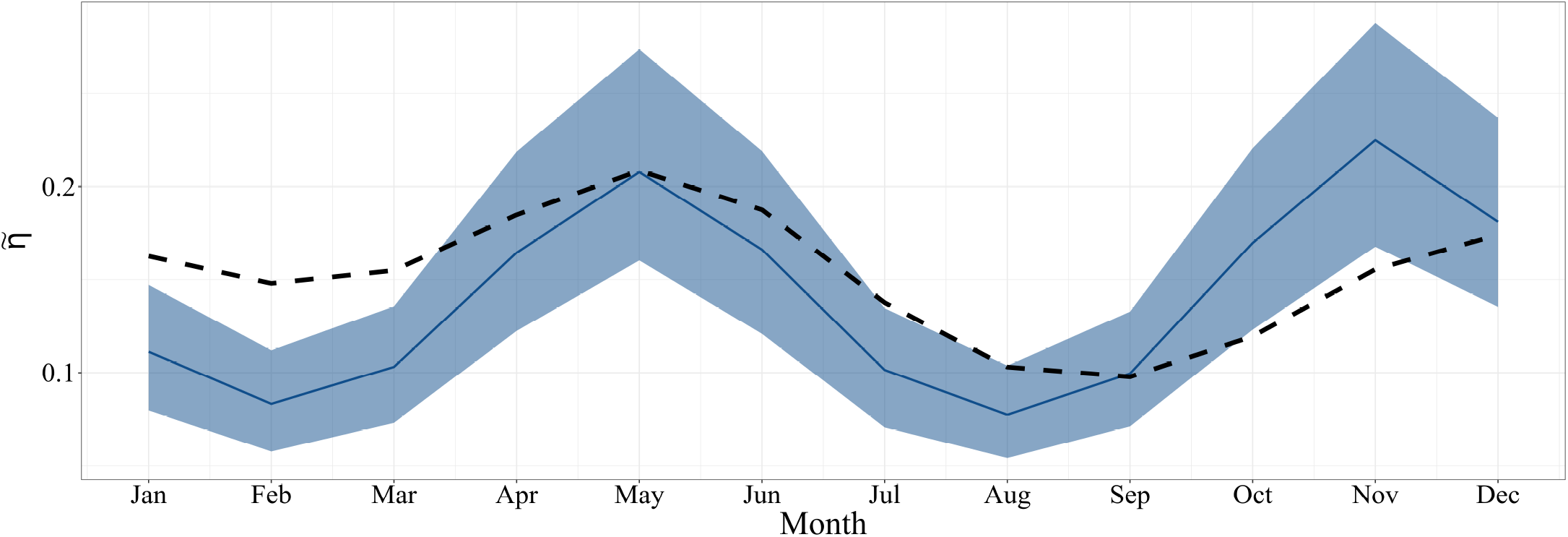
Conditional probabilities of encounter as a function of month. The solid line denotes the maximum *a posteriori* post-release survival estimate, whereas the band denotes the 80% credible interval. The dashed black line denotes the posterior median seasonal recreational angler reef fish effort for the West Florida Peninsula, scaled for comparability, from Hyman et al. (2024)

## References

Alonso-Fernández, A., G. Mucientes, and D. Villegas-Ríos (2022). “Discard survival of coastal elasmobranchs in a small-scale fishery using acoustic telemetry and recapture data”. In: Estuarine, Coastal and Shelf Science 276, p. 108037. doi: 10.1016/j.ecss.2022.108037.

Bacheler, N. M., J. A. Buckel, J. E. Hightower, L. M. Paramore, and K. H. Pollock (2009). “A combined telemetry–tag return approach to estimate fishing and natural mortality rates of an estuarine fish”. In: Canadian Journal of Fisheries and Aquatic Sciences 66.8, pp. 1230–1244. doi: 10.1139/F09-07.

Benoît, H. P., M. Morfin, and C. W. Capizzano (2020). “Improved estimation of discard mortality rates with in situ experiments involving electronic and traditional tagging”. In: Fisheries Research 221, p. 105398. doi: 10.1016/j.fishres.2019.105398.

Biesinger, Z., B. M. Bolker, D. Marcinek, and W. J. Lindberg (2013). “Gag (textitMycteroperca microlepis) space-use correlations with landscape structure and environmental conditions”. In: Journal of Experimental Marine Biology and Ecology 443, pp. 1–11. doi: 10.1016/j.jembe.2013.02.004.

Bohaboy, E. C., T. L. Guttridge, N. Hammerschlag, M. P. Van Zinnicq Bergmann, and W. F. Patterson III (2020). “Application of three-dimensional acoustic telemetry to assess the effects of rapid recompression on reef fish discard mortality”. In: ICES Journal of Marine Science 77.1, pp. 83–96. doi: 10.1093/icesjms/fsz202.

Bower, S. D., Ø. Aas, R. Arlinghaus, T. Douglas Beard, I. G. Cowx, A. J. Danylchuk, K. M. Freire, W. M. Potts, S. G. Sutton, and S. J. Cooke (2020). “Knowledge gaps and management priorities for recreational fisheries in the developing world”. In: Reviews in Fisheries Science & Aquaculture 28.4, pp. 518–535. doi: 10.1080/23308249.2020.1770689.

Brownscombe, J. W., K. Hyder, W. Potts, K. L. Wilson, K. L. Pope, A. J. Danylchuk, S. J. Cooke, A. Clarke, R. Arlinghaus, and J. R. Post (2019). “The future of recreational fisheries: advances in science, monitoring, management, and practice”. In: Fisheries Research 211, pp. 247–255. doi: 10.1016/j.fishres.2018.10.019.

Bürkner, P.-C. (2017). “brms: An R package for Bayesian multilevel models using Stan”. In: Journal of statistical software 80, pp. 1–28.

Burns, K. M., C. C. Koenig, and F. C. Coleman (2002). Evaluation of Multiple Factors Involved in Release Mortality of Undersized Red Grouper, Gag, Red Snapper, and Vermilion Snapper. Technical Report No. 790. Mote Marine Laboratory. url: https://sedarweb.org/documents/s24rd44-evaluation-of-multiple-factors-involved-in-release-mortality-of-undersized-red-grouper-gag-red-snapper-and-vermilion-snapper/.

Burns, K. M. (2009). “Differences between Red Grouper (Epinephelus morio) and Red Snapper (Lutjanus campechanus) Swim Bladder Morphology and How These Differences Affect Survival during Rapid Depressurization”. In: Evaluation of the Efficacy of the Minimum Size Rule in the Red Grouper and Red Snapper Fisheries With Respect to J and Circle Hook Mortality and Barotrauma and the Consequences for Survival and Movement. url: https://digitalcommons.usf.edu/etd/1881/.

Campbell, M. D., A. G. Pollack, W. B. Driggers, and E. R. Hoffmayer (2014). “Estimation of hook selectivity of Red Snapper and Vermilion Snapper from fishery independent surveys of natural reefs in the northern Gulf of Mexico”. In: Marine and Coastal Fisheries 6.1, pp. 260–273. doi: 10.1080/19425120.2014.968302.

Chih, C.-P. (2006). Estimation of species misidentification in the commercial landing data of gag groupers and black groupers in the South Atlantic (SEDAR10-DW-28). Tech. rep. SFD-2006-013. Miami, FL, USA: National Marine Fisheries Service, Southeast Fisheries Science Center, NOAA. url: https://sedarweb.org/documents/s10dw28-species-id-south-atlantic-eta-1-week-post-workshop/.

Christiansen, H. M., J. J. Solomon, T. S. Switzer, and R. B. Brodie (2022). “Assessing the size selectivity of capture gears for reef fishes using paired stereo-baited remote underwater video”. In: Fisheries Research 249, p. 106234. doi: 10.1016/j.fishres.2022.106234.

Christiansen, H. M., T. S. Switzer, S. F. Keenan, A. J. Tyler-Jedlund, and B. L. Winner (2020). “Assessing the relative selectivity of multiple sampling gears for managed reef fishes in the eastern Gulf of Mexico”. In: Marine and Coastal Fisheries 12.5, pp. 322–338. doi: 10.1002/mcf2.10129.

Coggins, L. G., M. J. Catalano, M. S. Allen, W. E. Pine III, and C. J. Walters (2007). “Effects of cryptic mortality and the hidden costs of using length limits in fishery management”. In: Fish and Fisheries 8.3, pp. 196–210. doi: 10.1111/j.1467-2679.2007.00247.x.

Curtis, J. M., M. W. Johnson, S. L. Diamond, and G. W. Stunz (2015). “Quantifying delayed mortality from barotrauma impairment in discarded red snapper using acoustic telemetry”. In: Marine and Coastal Fisheries 7.1, pp. 434– 449. doi: 10.1080/19425120.2015.1074968.

Ferter, K., M. S. Weltersbach, O.-B. Humborstad, P. G. Fjelldal, F. Sambraus, H. V. Strehlow, and J.H. Vølstad (2015). “Dive to survive: effects of capture depth on barotrauma and post-release survival of Atlantic cod (Gadus morhua) in recreational fisheries”. In: ICES Journal of Marine Science 72.8, pp. 2467–2481. doi: 10.1093/icesjms/fsv102.

Gelman, A., D. Lee, and J. Guo (2015). “Stan: a probabilistic programming language for Bayesian inference and optimization”. In: Journal of Educational and Behavioral Statistics 40.5, pp. 530–543. doi: 10.3102/1076998615606113.

Hueter, R. E., C. A. Manire, J. P. Tyminski, J. M. Hoenig, and D. A. Hepworth (2006). “Assessing mortality of released or discarded fish using a logistic model of relative survival derived from tagging data”. In: Transactions of the American Fisheries Society 135.2, pp. 500–508. doi: 10.1577/T05-065.1.

Hyder, K., M. S. Weltersbach, M. Armstrong, K. Ferter, B. Townhill, A. Ahvonen, R. Arlinghaus, A. Baikov, M. Bellanger, J. Birzaks, et al. (2018). “Recreational sea fishing in Europe in a global context—participation rates, fishing effort, expenditure, and implications for monitoring and assessment”. In: Fish and Fisheries 19.2, pp. 225–243. doi: 10.1111/faf.12251.

Hyman, A. C., C. Ramsay, T. A. Cross, B. Sauls, and T. K. Frazer (2026a). “Leveraging statistical models to improve preseason forecasting and in-season management of a recreational fishery”. In: North American Journal of Fisheries Management 46.1, pp. 222–236. doi: 10.1093/najfmt/vqaf100.

Hyman, A. C., C. Ramsay, S. Wilms, and T. K. Frazer (2026b). “Depth and water temperature drive elevated post-release mortality of gray triggerfish (Balistes capriscus)”. In: Frontiers in Marine Science 13, p. 1743605. doi: 10.3389/fmars.2026.1743605.

Hyman, A. C., D. Chagaris, M. Drexler, and T. K. Frazer (2024). “Modeling effort in a multispecies recreational fishery; Influence of species-specific temporal closures, relative abundance, and seasonality on monthly angler-trips”. In: Fisheries Research 279, p. 107136. issn: 0165-7836. doi: 10.1016/j.fishres.2024.107136.

Hyman, A. C., D. Chagaris, and T. K. Frazer (2025). “Influence of temporal regulations on harvest and discards in the recreational Gulf of Mexico gag (Mycteroperca microlepis) fishery”. In: Fisheries Research 285, p. 107332. issn: 01657836. doi: 10.1016/j.fishres.2025.107332.

Hyman, A. C., J. Cortes, L. Eguia, S. Wilms, and T. K. Frazer (2026c). “Depth and sea surface temperature drive post-release condition of reef-associated fishes”. In: Fisheries Research 296, p. 107700. issn: 0165-7836. doi: 10.1016/j.fishres.2026.107700.

Ihde, T. F., M. J. Wilberg, D. A. Loewensteiner, D. H. Secor, and T. J. Miller (2011). “The increasing importance of marine recreational fishing in the US: challenges for manage-ment”. In: Fisheries Research 108.2-3, pp. 268–276. doi: 10.1016/j.fishres.2010.12.016.

Lindberg, W. J., T. K. Frazer, K. M. Portier, F. Vose, J. Loftin, D. J. Murie, D. M. Mason, B. Nagy, and M. K. Hart (2006). “Density-dependent habitat selection and performance by a large mobile reef fish”. In: Ecological Applications 16.2, pp. 731–746. doi: 10.1890/1051-0761(2006)016[0731:DHSAPB]2.0.CO;2.

McGovern, J. C., G. R. Sedberry, H. S. Meister, T. M. Westendorff, D. M. Wyanski, and P. J. Harris (2005). “A tag and recapture study of gag, Mycteroperca microlepis, off the southeastern US”. In: Bulletin of Marine Science 76.1, pp. 47–59.

Pollock, K. H., H. Jiang, and J. E. Hightower (2004). “Combining telemetry and fisheries tagging models to estimate fishing and natural mortality rates”. In: Transactions of the American Fisheries Society 133.3, pp. 639–648. doi: 10.1577/T03-029.1.

Ramsay, C., M. D. Campbell, and B. Sauls (2025). “A meta-analysis of immediate and delayed discard mortality of red snapper (Lutjanus campechanus)”. In: Fisheries Research 291, p. 107524. doi: 10.1016/j.fishres.2025.107524.

Rudershausen, P. J., J. A. Buckel, and J. E. Hightower (2014). “Estimating reef fish discard mortality using surface and bottom tagging: effects of hook injury and barotrauma”. In: Canadian Journal of Fisheries and Aquatic Sciences 71.4, pp. 514–520. doi: 10.1139/cjfas-2013-0337.

Rudershausen, P. J., S. J. Poland, W. Merten, and J. A. Buckel (2019). “Estimating Discard Mortality for Dolphin-fish in a Recreational Hook-and-Line Fishery”. In: North American Journal of Fisheries Management 39.6, pp. 1143– 1154. doi: 10.1002/nafm.10348.

Rudershausen, P. J., B. J. Runde, R. M. Tharp, J. H. Merrell, N. M. Bacheler, W. F. Patterson III, and J. A. Buckel (2025). “Discard mortality rates of Red Snapper after barotrauma and hook trauma: Insights from using acoustic telemetry in the US South Atlantic”. In: North American Journal of Fisheries Management, vqaf012. doi: 10.1093/najfmt/vqaf012.

Rudershausen, P. J., H. M. Schmidt, J. H. Merrell, B. J. Runde, and J. A. Buckel (2023). “Effectiveness of venting and recompression for increasing postrelease survival of barotraumatized Black Sea Bass across a range of depths”. In: North American Journal of Fisheries Management 43.1, pp. 257–267. doi: 10.1002/nafm.10864.

Rudershausen, P., B. Runde, and J. Buckel (2020). “Effectiveness of venting and descender devices at increasing rates of postrelease survival of Black Sea Bass”. In: North American Journal of Fisheries Management 40.1, pp. 125–132. doi: 10.1002/nafm.10387.

Runde, B. J., N. M. Bacheler, K. W. Shertzer, P. J. Rudershausen, B. Sauls, and J. A. Buckel (2021). “Discard mortality of Red Snapper released with descender devices in the US South Atlantic”. In: Marine and Coastal Fisheries 13.5, pp. 478–495. doi: 10.1002/mcf2.10175.

Runde, B. J., P. J. Rudershausen, B. Sauls, C. S. Mikles, and J. A. Buckel (2019). “Low discard survival of gray triggerfish in the southeastern US hook-and-line fishery”. In: Fisheries Research 219, p. 105313. doi: 10.1016/j.fishres.2019.105313.

Sauls, B. (2014). “Relative survival of gags Mycteroperca microlepis released within a recreational hook-and-line fishery: Application of the Cox Regression Model to control for heterogeneity in a large-scale mark-recapture study”. In: Fisheries Research 150, pp. 18–27. doi: 10.1016/j.fishres.2013.10.008.

Scheffel, T. K., J. E. Hightower, J. A. Buckel, J. R. Krause, and F. S. Scharf (2020). “Coupling acoustic tracking with conventional tag returns to estimate mortality for a coastal flatfish with high rates of emigration”. In: Canadian Journal of Fisheries and Aquatic Sciences 77.1, pp. 1–22. doi: 10.1139/cjfas-2018-0174.

SEDAR72 (2021). Stock Assessment Report: Gulf of Mexico Gag Grouper. Tech. rep. 4055 Faber Place Drive, Suite 201 North Charleston, SC 29405, USA: Southeast Data, Assessment, and Review. url: https://sedarweb.org/assessments/sedar-72/.

Shertzer, K., S. Crosson, E. Williams, J. Cao, R. DeVictor, C. Dumas, and G. Nesslage (2024). “Fishery management strategies for Red Snapper in the southeastern US Atlantic: A spatial population model to compare approaches”. In: North American Journal of Fisheries Management 44.1, pp. 113–131. doi: 10.1002/nafm.10966.

Shertzer, K. W., E. H. Williams, J. K. Craig, E. E. Fitzpatrick, N. Klibansky, and K. I. Siegfried (2019). “Recreational sector is the dominant source of fishing mortality for oceanic fishes in the Southeast United States Atlantic Ocean”. In: Fisheries Management and Ecology 26.6, pp. 621–629. doi: 10.1111/fme.12371.

Sivula, T., M. Magnusson, A. A. Matamoros, and A. Vehtari (2020). “Uncertainty in Bayesian leave-one-out crossvalidation based model comparison”. In: arXiv preprint 2008.10296. doi: 10.48550/arXiv.2008.10296.

Stallings, C. D., O. Ayala, T. A. Cross, and B. Sauls (2023). “Post-release survival of red snapper (Lutjanus campechanus) and red grouper (Epinephelus morio) using different barotrauma mitigation methods”. In: Fisheries Research 264, p. 106717.

Stan Development Team (2022). The Stan Core Library. Version 2.32. url: http://mc-stan.org/%2011.

Tetzlaff, J. C., W. E. Pine, M. S. Allen, and R. N. Ahrens (2013). “Effectiveness of size limits and bag limits for managing recreational fisheries: a case study of the Gulf of Mexico recreational gag fishery”. In: Bulletin of Marine Science 89.2, pp. 483–502. doi: 10.5343/bms.2012.1025.

Trudeau, A., E. A. Bochenek, A. S. Golden, M. C. Melnychuk, D. R. Zemeckis, and O. P. Jensen (2022). “Lower possession limits and shorter seasons directly reduce for-hire fishing effort in a multispecies marine recreational fishery”. In: Canadian Journal of Fisheries and Aquatic Sciences 79.8, pp. 1211–1224. doi: 10.1139/cjfas-2021-0137.

Vehtari, A., J. Gabry, M. Magnusson, Y. Yao, P.-C. Bürkner, T. Paananen, and A. Gelman (2022). loo: Efficient leaveone-out cross-validation and WAIC for Bayesian models. R package version 2.5.1. url: https://mc-stan.org/loo/.

Vehtari, A., A. Gelman, and J. Gabry (2017). “Practical Bayesian model evaluation using leave-one-out crossvalidation and WAIC”. In: Statistics and Computing 27.5, pp. 1413–1432. doi: 10.1007/s11222-016-9696-4.

Williams-Grove, L. J. and S. T. Szedlmayer (2016). “Mortality estimates for red snapper based on ultrasonic telemetry in the Northern Gulf of Mexico”. In: North American Journal of Fisheries Management 36.5, pp. 1036–1044. doi: 10.1080/02755947.2016.1184197.

Zimmermann, S., L. Kehoe, M. Taylor, J. H. Tarnecki, N. Bacheler, Z. A. Siders, and W. F. Patterson (2025). “Post-release Mortality of Red Snapper, Lutjanus campechanus, in US Atlantic Waters off Northeast Florida Estimated with Three-Dimensional Acoustic Telemetry”. In: SSRN Electronic Journal. doi: 10.2139/ssrn.5430934. url: https://ssrn.com/abstract=5430934.

